# Immune and non-immune cell fencing of tumor cells is a widespread and functionally relevant spatial pattern in solid cancers

**DOI:** 10.1101/2025.08.11.669531

**Authors:** Giuseppe Giuliani, Dhruv Chavan, Rushil Gaddam, William C. Stewart, Zihai Li, Ciriyam Jayaprakash, Jayajit Das

## Abstract

Solid tumors are characterized by a spatially heterogeneous mixture of cancer cells, immune cells and other non-tumor cells. Recent characterization of the heterogeneity at the single cell level has revealed spatial patterns of different cell types often lacking a simple geometric structure associated with cancer progression. Here we investigated the occurrence of physical fencing of tumor cells by specific immune and non-immune phenotypes in the tumor microenvironment (TME) and the association of these clusters to cancer progression in a wide range solid tumors formed in different organs. We analyzed published patient response and imaging mass cytometry (IMC) datasets from tumor microarrays obtained from tumor tissues in triple-negative breast cancer (279 patients), lung cancer (416 patients), melanoma (30 patients), colorectal cancer (9 patients), glioma (185 patients), and head and neck cancer (139 patients) to characterize the presence of fencing clusters of various cell types and their association with differing patient outcomes. Devising and employing simple mechanistic and stochastic spatially-resolved computational models we quantify the dependence of the pro- and anti-tumor roles of a fencing cluster on the size and the lifetime of the cluster, as well as the chemokine gradient in the local environment. We unveiled that spatial patterns of immune cells, especially through fencing tumor boundary, affects tumor progression and treatment responsiveness to immunotherapy.

## Introduction

A tumor is a spatially heterogeneous tissue composed of tumor cells, various immune cells, connective tissues, blood and lymphatic vessels, and extracellular materials like collagen. Spatial profiling of solid tumors using proteomics and transcriptomics experiments show wide variations of spatial co-organization of tumor and immune cells across length scales of 10μm to over several millimeters spanning the entire tumor(1). The spatial organization of tumor and immune cells in primary tumors originating in the same organ also varies widely across patients. Our understanding of the spatial heterogeneity of tumors and its role in tumor progression remains limited and is only beginning to emerge(1-3). Some spatial features such as tertiary lymphoid structures (TLS) where T- and B-cells co-cluster in length scales of 50-200μm have been found in many solid cancers including breast, lung, and skin cancer and have been associated with a positive patient prognosis(4). Several recent studies have also found increased infiltration of lymphocytes within the tumor (‘inflamed’ immunophenotype) in breast cancer as opposed to lymphocytes mostly residing at the tumor boundary (‘excluded’ immunophenotype) is associated with favorable response to immunotherapy in the patients(5). Application of machine learning approaches found spatial regions lacking simple geometric structures enriched in specific subtypes of immune cells associated with functional consequences for cancer prognosis(6-9). Thus, it appears there are spatial features that can persist across heterogeneities and occur in patient cohorts associated with different disease outcomes. Some of these spatial signatures may even be potential spatial biomarker candidates. However, mechanistic roles of these spatial patterns to regulate tumor growth and patient outcome remain poorly characterized.

Tumor growth is shaped by spatially localized interactions between immune and tumor cells. Given the spatial heterogeneity of the tumor microenvironment (TME) some of the local organization of tumor and immune cells provide niches for tumor growth and generate similar phenotypic response(2, 6, 10). Spatial profiling of tumors is typically done with resected or biopsied tumor tissues and thus provides a time-stamped snapshot of the dynamic tumor progression. Therefore, a critical gap exists in mechanistically relating a specific spatial pattern of tumor and immune cells gleaned from spatial profiling data to tumor growth and eventually to patient response. Mechanistic computational models in particular agent-based models (ABMs) where agents describe individual components of the TME such as tumor and immune cells which interact via specific rules have been employed to bridge the gap(3, 11, 12). A recent ABM trained on multiplex imaging data from melanoma tumor tissues in mice described interaction of different phenotypes (activated/exhausted) of T cells with tumor cells and found that the initial spatial distribution of T cells relative to each other and cancer cells may enhance or limit the exhaustion of T cells(11).

Interactions between immune and tumor cells at the tumor immune boundary have been found to be important for tumor growth(13, 14). Investigation of tumor immune boundaries using imaging mass cytometry (IMC) of breast tumor microarrays (TMA) of sizes ≤ 1 mm assaying over 30 different protein markers in single cells reveal more transcriptionally active tumor cells at the boundary compared to the tumor core (13). In our earlier analysis and agent-based modeling of IMC data obtained from patient melanoma, TMAs of sizes ≤ 1 mm showed ‘fencing’ of tumor cells by exhausted CD8+ T or T_ex_ cells in our model simulations(3). This fencing was also observed in single cell CyCIF imaging of melanoma tumors in patients (14, 15). In our fencing pattern, tumor cells are surrounded by clusters of T_ex_ cells. The T_ex_ cells potentially generate a barrier for immune cells such as cytotoxic CD8+ T cells that can lyse the tumor thus playing a pro-tumor role. In contrast, fencing of tumor cells by activated CD8+ T cells could lead to an increased rate of lysis of the tumor cells. Thus, fencing of tumor cells by immune cells that are likely to be present at the tumor boundaries could have different functional responses depending on the immune cell type and the context.

Here we investigated if fencing of tumor cells by different immune cells in different solid cancers occurs in patient cohorts and are associated with survival or patient response to immunotherapy. We analyzed published IMC datasets of TMAs obtained from six different solid cancers (skin, colorectal, lung, brain, breast, and head and neck) to determine the presence of fencing structures composed of different immune and non-immune cells and then evaluated their association with patient response and survival. We observed with statistical significance that fences composed of immune cells such as T, myeloid and NK cells, and non-immune cells such as fibroblasts are present in the solid tumors examined here. In addition, we found statistically significant associations of such fencing structures with functional outcomes in patients such as response to immune checkpoint therapy and survivability. Finally, we simulated a spatially-resolved mechanistic ABM to theoretically evaluate functional implications of the fencing formation. We found that in silico, through different mechanisms, cell fences can diminish or augment the access of immune cells to cancer cells and, in one case, limit their control of the cancer cell population.

## Results

### 1. Fencing of tumor cells by immune and non-immune cells is observed in solid tumors of various organs

We investigated the existence of fencing structures formed by non-tumor cells in the tumor microenvironment as defined below. We represented the single cells in the IMC dataset as nodes of a graph where we connected a pair of single cells (or nodes) by an edge if the spatial distance between the pair is ≤ 30 μm. We defined a spatial cell cluster as a group of three or more immune or non-immune cells of the same phenotype (e.g. CD8+ T cell) connected by a continuous path of edges (**Figure 1a**). A fencing cluster is then a spatial cluster with at least one of the (non-tumor) cells is edge connected with a tumor cell.

**Figure 1.**
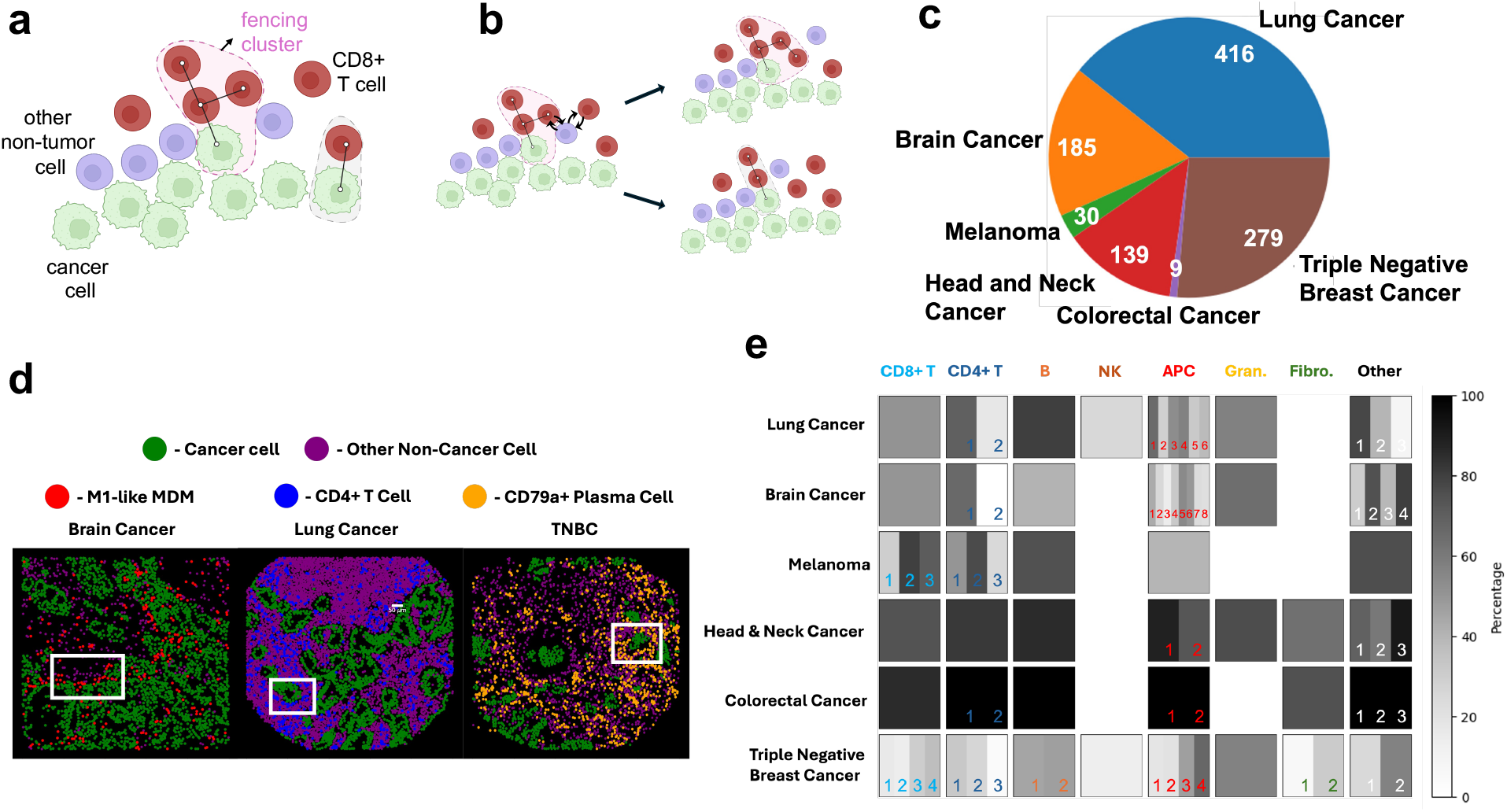
Fencing of tumor cells by immune and non-immune cells is found in solid tumors of various organs. **(a)** Defining fencing clusters composed of CD8+ T cells on the tumor periphery. Cell positions in the patient slide data are provided by the centroid of the cell nucleus. Two cells are in contact if their centroids are within 30 *µ* of each other and are connected by black edges. When at least three cell nodes of the same type are connected by a continuous path of edges, they define a cluster. If at least one of the cells in a cluster is in contact with at least one cancer cell, the cluster is a fencing cluster. **(b)** A schematic displaying how non-cancer cells are permuted to build new randomized configurations of a patient slide. This basic permutation generates randomized configurations discussed in Subsection 1 of Results. **(c)** The number of cancer patients from which IMC slides were produced of their tumor for each solid cancer type. Clockwise, the cancer types are lung (adenocarcinoma), triple negative breast cancer (TNBC), colorectal cancer, head and neck cancer (HNC), melanoma, and brain cancer (glioma and brain metastases) **(d)** Snapshots of the TME generated from IMC data from patients with fencing of three different cell phenotypes in three different solid cancer types: M1-like monocyte-derived macrophages (MDMs) in a metastasis to the brain, CD4+ T cells in lung cancer and CD79a+ plasma cells in triple-negative breast cancer (TNBC). Each slide displays points representing the centroid of each cell in the system. The other non-cancer cells in purple refers to phenotypes that are not part of the fencing. The white boxes outline one region in each slide where the chosen cell phenotype forms fencing clusters on the boundary of the cancer cells. **(e)** A table showing the fraction of slides which display fencing significantly more than random for each non-cancer cell phenotype in each cancer dataset. Each box falls within a column designating a general cell grouping (e.g., CD8+ T, APC) and a row designating the solid cancer type. Within each tumor type and cell group combination, there may be 0 (white spaces with no box outline) or more cell types depending on the dataset. Each subregion of a box represents an individual cell phenotype within the cell group. The shade of the subregion, in reference to the color bar (right), demarks the percentage of slides which display significant (controlled at p<0.05) fencing of that specific cell phenotype. Boxes with more than one cell phenotype have numbers which may be used as reference when reading **table S1**, which explicitly states each cell phenotype and the corresponding percentage of slides which display statistically significant fencing. Tregs (in positions 2, 2, 3, 2, and 3 in lung, brain, melanoma, colorectal and breast cancers respectively) only form fencing clusters very rarely except for in colorectal cancer. See subsection 1 of Results for a more detailed discussion.

We analyzed IMC datasets obtained from TMAs for a wide variety of solid cancers (**Figure 1c**) to determine how often different cell types give rise to fencing clusters across patients in a statistically significant manner. The datasets considered span triple-negative breast cancer (TNBC) (8), lung adenocarcinoma cancer (6), melanoma (16), colorectal cancer (CRC) (17), glioma (primary glioma and metastases to the brain) (7) and head and neck cancer (HNC) (18) (**Figure 1c**). For each solid cancer, we analyzed IMC data yielded from patient TME slides to identify fencing clusters for a variety of immune and non-immune cells (**Figure 1d**). To determine if these clusters could have occurred by chance we proceeded as follows. For each cell phenotype *g* (e.g., CD8+ T cell), we calculated the probability that the *g*-cell fencing clusters found in the data could occur in a random spatial distribution of all the non-tumor cells (including *g-*cells) present in the TMA. To do this, we generated an ensemble of spatial configurations by randomly exchanging the cell phenotype of all *g*-cells with any of the other non-tumor single cells (including other *g-*cells).

The permuted configuration preserves the spatial structure of the tumor cells and non-tumor cells but randomly shuffles the positions of all individual *g*-cells (a subset of which form fencing clusters) into the population of non-tumor cells. This can lead to fragmentation, dissolution, or enlargement of the original fencing clusters (**Figure 1b**). We generated over 1000 permuted configurations for each TMA slide and evaluated the probability of there being at least as many *g-*cells observed in fencing clusters in the original configuration given the spatial distribution of the non-tumor and tumor cells in the original slide. This probability is a p-value for the occurrence of the observed fencing clusters in each slide (see Materials and Methods for details). We control for a p-value ≤ 0.05 to determine if a patient slide contains fencing clusters formed by a cell phenotype *g*. We finally obtain the fraction (*f*_*p*_) of the patient slides in which the fencing clusters of cell phenotype *g* occur significantly.

We summarize the results for the frequency of occurrence of different cell types in the fencing clusters seen in various cancers (**Figure 1e, Table S1**). Among immune cells, we find a CD8+ and CD4+ T cells and different subtypes of the T cells, such as regulatory CD4+ T cells (CD4+ FOXP3+), activated (CD8+GZMB +) and exhausted (CD8+ PD1+) CD8+ T cells form fencing structures in all the solid tumors investigated here. Almost all the patient slides in melanoma, HNC, and colorectal cancers show fencing clusters formed by CD8+/CD4+ T cells and their subtypes. The TMAs for the lung, glioma, and the breast cancers show a lower occurrence frequency of T cell (CD4+/CD8+, not Tregs) fencing with < 70% patients as compared to melanoma, HNC and CRC. Fencing of CD4+ T cells (not including Tregs) occurred more often in patients as compared to CD8+ T cell fencing in lung, glioma, HNC, and CRC patients. Regulatory T cell (Treg) fencing clusters are less likely to form fencing clusters as Treg fencing clusters are only found in ≤ 25% of patients with the exception of in CRC. The lymphocyte B cells gave rise to fencing clusters in a large fraction (≥75%) of the patients for the lung cancer, melanoma, HNC, and CRC and in a smaller fraction (<48%) patients for glioma and TNBC. NK cells were the least likely to give rise to fencing clusters, present only in a small fraction of patients (<26%) in lung and TNBC.

### 2. Fencing of tumor cells by immune and non-immune cells is associated with patient response

We investigated if increased participation of a non-tumor cell type in fencing clusters is associated with patient clinical outcomes such as response to cancer therapy, patient survival or cancer progression. We used published IMC data for TMAs obtained from solid tumors in patients (**Figure 1c,d**) accompanied by the corresponding clinical outcomes and Kaplan-Meier survival data for those patients. Cancer therapy in these patients ranged from immune checkpoint blockage (ICB) therapy, chemotherapy, combination of chemotherapy and ICB. In addition, for brain cancer, we also investigated if the increased participation of non-tumor cells in the fencing clusters is associated with the origin of the tumor, namely primary origin (e.g., glioblastoma) or brain metastasis. We quantified the increased participation of non-tumor cells in fencing clusters in the following way.

The fencing participation metric is defined to capture the propensity of *g-*cells to occur in fencing clusters in a patient slide as compared to when the cells in the patient slide are permuted randomly as described. The fencing participation metric (FPM) is defined by 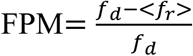 where the numerator is the difference between the number of *g*-cells in fencing structures in a patient slide, *f*_*d*_, and the ensemble average number of *g*-cells in fencing structures across the randomized cell configurations of the slide or < *f*_*r*_ >. The normalization by *f*_*d*_ is chosen so that the metric is one when a patient shows *g*-cell fencing structures, but the randomized slides show no *g*-cell fencing structures. It decreases to zero when the average number of *g-*cells in the fencing structures in the randomized slides increases to the be the same or a larger than the number found in the original patient slide. We mapped negative values of FPM to zero, thus, when more cells are found in the fencing structures in the randomized slides, we denote those as no more cells in fencing structures found in a patient slide as compared to the randomized slides. We evaluate the FPM for each fencing cell phenotype *g* for patient slides with at least 50 *g*-cells and 50 cancer cells. Comparing the distributions of the FPM between patient clinical outcomes, such as response or no response to therapy, yields significant relationships between the FPM and patient tumor progression (See Materials and Methods for more details). As the FPM of a given cell type depends on the spatial distribution of cells in the TME, it innately has a dependence on that cell’s density. We checked that the correlations that the FPM has with patient clinical outcome are not solely due to its dependencies on cell densities rather than more complex spatial relationships. We confirmed that the FPM not only provides additional information about patient clinical outcome as compared to cell densities but can have a stronger correlation with patient clinical outcome than cell densities (See Materials and Methods for more details).

We compared the distributions of the FPM evaluated across the patient slides for various cell types between patient clinical outcomes to find statistically significant differences. In HNC, the average FPM values for adaptive lymphocytes such as CD8+ or CD4+ T cells, and B cells, and myeloid cells such as macrophages and granulocytes show higher magnitudes in responders to therapy compared to non-responders (**Figure 2a**). Using a similar comparison, we found increased participation in fencing clusters by non-immune cells, such as fibroblasts and cells forming lymph vessels, is associated with responders as opposed to non-responders to therapy. Next, we compared the association of the survivability of the patients with FPM for the immune and non-immune cells. We found increased values of FPM are associated with improved survivability in the HNC patients (**Figure 2d-f**). In TNBC, responders to therapy exhibit lower average FPMs for GZMB+ CD8+ T cell, PD1+ CD4+ T cell, Treg, CD79a+ plasma cell, PDL1+ APC and myofibroblast (**Figure 2b**). In contrast, the average FPM for TCF1+ CD4+ T cells and neutrophils in responder slides are larger as compared to non-responders to therapy. This points to fences mechanistically playing pro-as well as anti-tumor roles which we will investigate in the next section. In brain cancer, we found that the average FPM of CD4+ T cells, M1-like monocyte-derived macrophages (MDM) and other (not CD8+ or CD4+) T cells is higher in primary gliomas as compared to in metastases to the brain. The opposite is true for non-classical (non-cl.) monocytes and M1-like microglia (**Figure 2c**). Patients with lower endothelial cell FPM are associated with improved survivability (**Figure 2g**). We found no statistically significant associations (p≤ 0.05) of the FPM with patient response to therapy, tumor progression, or survivability in lung cancer, melanoma, and colorectal cancers. The melanoma and colorectal cancer datasets had a small number of patients (30 and 9 respectively), which could contribute to the absence of any statistically significant association with the patient response or metastatic status. In summary, these correlations imply that the formation of fencing structures from various cell types could influence tumor progression. Next, we consider theoretical models to elucidate possible effects of fencing on the progression of the TME.

**Figure 2.**
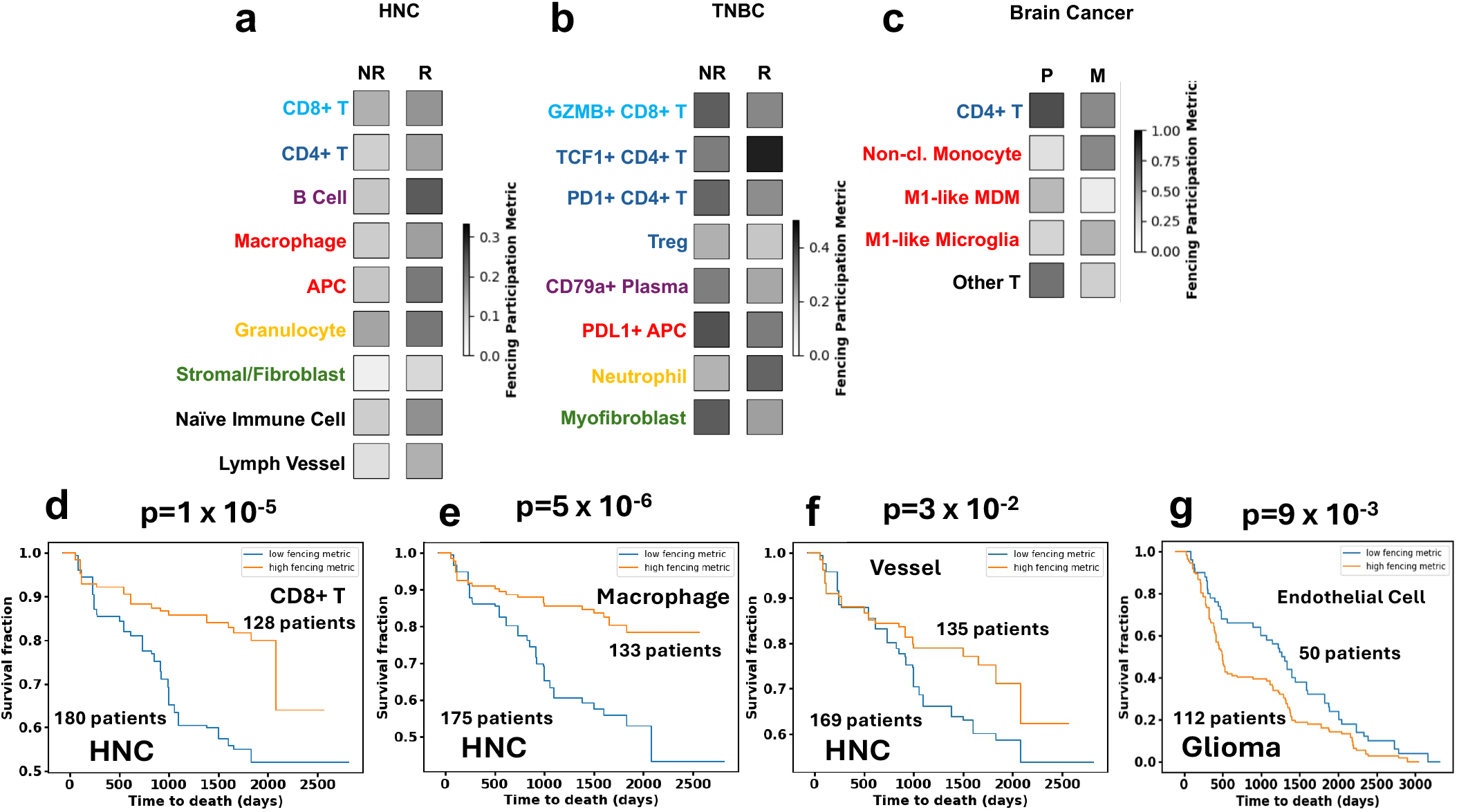
Fences formed by immune and non-immune cells are associated with patient responses. We compared the distributions of the fencing participation metric of any given cell type between responders (**R**) and non-responders (**NR**) to therapy for **(a)** HNC, **(b)** TNBC, and between metastatic tumors **(M**) to primary gliomas (**P**) for **(c)** brain cancer. In particular, we display the average fencing participation metrics (shaded with reference to scale bars) across responses for cell types whose average fencing participation metrics were found to differ significantly (p<0.05). Note that each cancer type has its own scale bar. In HNC, responders showed an increased average fencing participation across many cell types. In TNBC, we particularly found that a higher average fencing participation metric of TCF1+ CD4+ T cells in responders which may point to a pro-immune role for fencing clusters. We also observed that non-responders have a higher average PDL1+ APC fencing participation metric than responders maybe due to a pro-tumoral role of fencing clusters. Finally in brain cancer, we observed that primary gliomas have higher CD4+ T cell and M1-like MDM fencing participation metrics on average than in metastases to the brain. **(d-f)** For HNC, we found that the survival curves of patients with high (above average) and low (below average) fencing participation metric calculated for certain cell types differ significantly (using the log-rank test, p<0.05). Here we showed the survival curves for those cell types, the p-values separating the curves and the number of patients which contribute to each curve. Higher FPM across all of these cell type corresponds to better survival outcomes. **(g)** In primary glioma, the survival curves of patients with high (above average) and low (below average) endothelial cell fencing participation metric differ significantly. Patients with a lower endothelial cell FPM are associated with better survival.

The key takeaway points: 1) The formation of fencing clusters by non-tumor cells are associated with patient response. However, the underlying mechanisms (e.g., causative vs associative) remain speculative. 2) The fencing clusters in brain cancers appear to be associated with the primary origin of the tumor which could point the role of the differences in primary and metastatic tumors such as neoantigen load, tumor mutation burden. We discuss some of these points in the Discussion section. In this section, we found that the presence or lack of fences composed of specific cell phenotypes is associated with different clinical outcomes such as response to therapy. In the next section, we employ minimal spatially resolved mechanistic models to quantitatively investigate two potential specific pro- and anti-tumor roles of fencing clusters.

### 3. Fencing of tumor cells by immune and non-immune cells could positively or negatively impact tumor progression

Our investigations with IMC data obtained from TMAs in different solid cancers found that fencing clusters are present in many solid cancers and in many cases are associated with response to therapy and patient survivability. Given the experimental evidence for fences of different phenotypes being associated with differing clinical outcomes (e.g. response and non-response to therapy) in certain cancers, in this section we investigate two simple computational models that illustrate pro- and anti-tumor effects of fencing quantitatively. Due to the lack of detailed, dynamical, in vivo data, we built and studied simple computational models to uncover how fencing clusters affect the progression of the TME.

We employed a stochastic agent-based model (ABM), interacting cell system (ICS) model, that we developed previously in the context of melanoma (3) to test the effects fences have an effect on cancer cell population. The ICS model simulates stochastic time evolution of immune and tumor cells in the TMA that have been studied by IMC by modeling cell motility, cancer proliferation and cell-cell interactions such as activated CD8+ T cell exhaustion, cancer lysis and activated CD8+ T cell proliferation (**Figure 3a**) (See Materials and Methods for more details).

**Figure 3.**
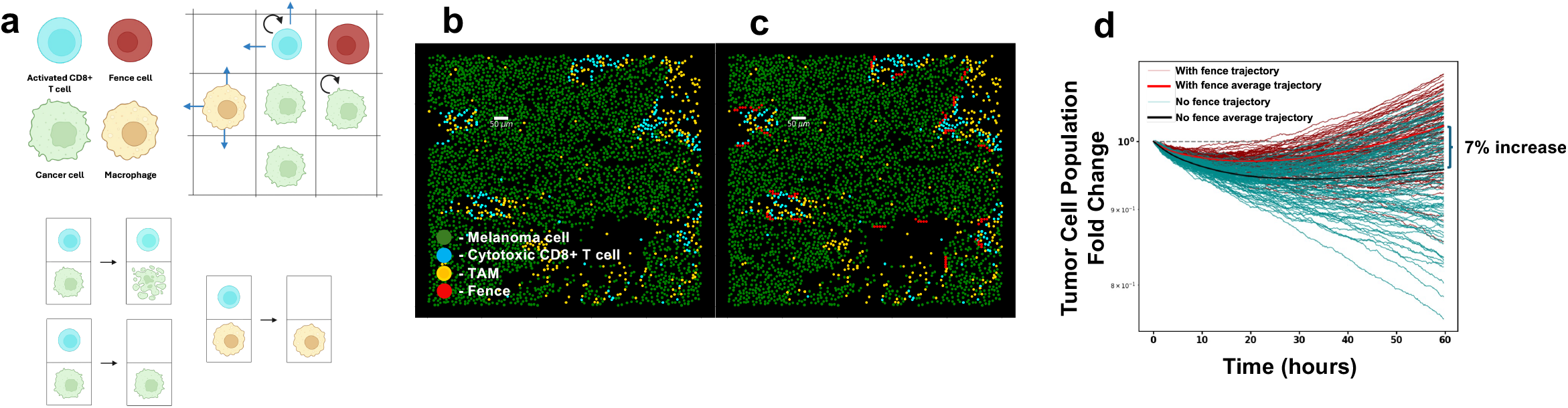
A stochastic ABM for melanoma characterizes the pro-tumor role of fencing. **(a)** The ABM consists of activated CD8+ T cells, non-tumor cells in the implanted fences, tumor associated macrophages (TAMs) and cancer cells interacting on a lattice. The cells are initially positioned (except the fences) as they are in patient data. The activated CD8+ T cells and TAMs move on the lattice but their movement is spatially restricted by the other cells on the lattice (including the cells constituting the fences). Cancer cells and activated CD8+ T cells proliferate. Some of the cell-cell interactions include cancer cell lysis and activated CD8+ T cell exhaustion by either cancer cells or TAMs. When activated CD8+ T cells are exhausted, they are removed from the system. **(b)** The original slide 06RD corresponding to a patient with melanoma who responded to ICI therapy. **(c)** The same patient slide with implanted fences of comparable size and density as those found in our analysis of TNBC. **(d)** Cancer cell population trajectories through time (2.5 days) when simulating the progression of the slides with (c) and without (b) the added fences with the modified ICS model (Results 3). Each initial condition is simulated 300 times. Here we plot a subset of the trajectories (100/300) in the interest of clarity. As the model is stochastic, there is variation in the outcome for each trajectory. The final ensemble average cancer cell population for the system with the fences is 7% larger than that for the original slide.

We studied the effect of fixed, impenetrable fences, roughly matching the average size (∼50 *µm* diameter) and density of non-cytotoxic fences (20 *mm*^−2^) found across the datasets, implanted into a patient slide by hand (**Figure 3b-c**). We first modified our earlier ICS model by removing exhausted CD8+ T cells to aid in the interpretation of results by avoiding the confounding effects of fences formed dynamically by T_ex_ cells (as shown in the earlier referenced work). Two cases were studied to investigate the effect of the fence: (1) the control, where the initial configuration of cells are obtained from an IMC slide corresponding to a responder patient to immunotherapy and (2) <u>the fence alternative</u>, where we modified the IMC slide from case #1 by implanting fences of non-tumor cells as described above.

As our model includes stochastic fluctuations in the processes considered, the kinetics of the tumor and non-tumor cells vary across different simulation runs even when started from the same initial configuration of cells. Therefore, to obtain the mean kinetics of the tumor and non-tumor populations, we average the populations across 300 simulation runs. The progression of average tumor and non-tumor cells over 2.5 days in the control and fence alternative were compared (**Figure 3d)**. We found the average of the cancer cell population at 2.5 days is 7% higher than that of the control case. The reduction of the access of activated CD8+ T cell to cancer cells due to the implanted fences resulted in an increase in cancer cell population growth.

To further clarify the effect of a fencing clusters in reducing the access of motile immune cells to cancer cells, we constructed a minimal two-dimensional spatially-resolved diffusion model **(Figure 4a**) which can be analyzed analytically or numerically. The minimal model considers a two-dimensional grid where a linear fence formed by a specific phenotype of immobile non-tumor cells resides on the tumor boundary represented by the bottom edge of the two-dimensional grid (**Figure 4a**) A different type of non-tumor cell which is motile is initially placed in the two-dimensional region. The motile non-tumor cell moves diffusively in the region. Once the motile cell reaches the tumor boundary it is taken out of the model, i.e., we impose an absorbing boundary condition. The fence acts as a barrier reflecting the diffusing cell thus blocking access to the tumor boundary. In order to isolate the effect of the fence on the accessibility of the cancer to non-tumor cells, we ignored cell proliferation and lysis in the model. Since the movement of the motile cell is stochastic in nature, the time the cell spends to access the tumor boundary starting from the same initial position can vary for different realizations of the kinetics. To quantify the effect of the fence in hindering the access of the motile cell to the tumor boundary, we computed the distribution of the time the motile non-tumor cell spends before reaching the tumor boundary across many realizations of the kinetics. This time distribution is also known as a first-passage time (FPT) distribution in probability theory. We found that, when a fixed fence is introduced, the FPT distribution becomes substantially broader compared to when the fence is absent, increasing the average time the motile cell takes to reach the tumor boundary (**Figure 4b**). This broadening is partly brought about by the change in distance between the initial position and the closest accessible portion of the tumor boundary (**Figure 4b**). However, the above FPT distribution depends on the initial position of the motile cell, the total time duration *T* over which the kinetics are studied, and the length of the fence. These variables are also relevant in the TME, as the motile cell (e.g., CD8+ T cell) can start in different positions, the size of the fence can vary, and fence might dissociate after a finite time τ_fence_. To account for the above dependencies, we average over the kinetics of motile non-tumor cells starting from positions throughout the system (**Figure 4a**). We defined the tumor access ratio as the ratio comparing the number of the diffusing cells which reach the tumor boundary within a time duration *T* when a fence of length *b* is present to those that reach the tumor boundary when the fence absent, i.e.,

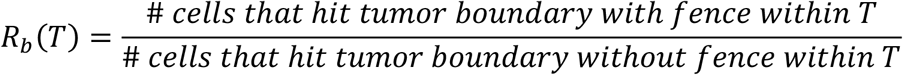

**Figure 4.**
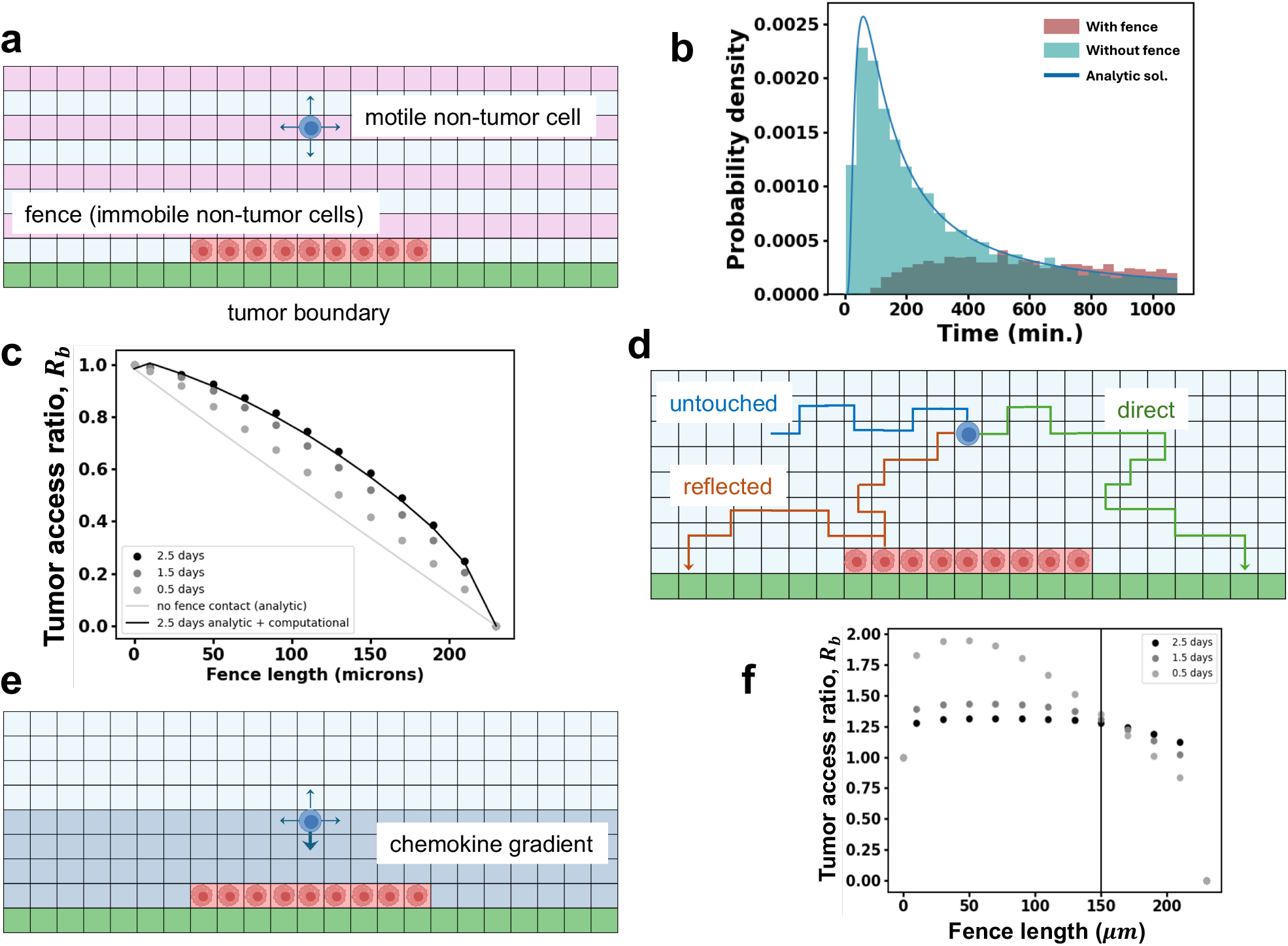
Stochastic and sptial minimal models quantify anti- and pro-tumor roles of fencing. **(a)** The initial spatial distribution of the simple model with periodic boundary conditions along the horizontal direction described in Materials and Methods. The green pixels represent the tumor boundary, the pink pixels represent starting positions, and the red pixels the cells in a fence of length 90 *µm* fence in a system of width 230 *µm* **(b)** The first-passage time (FPT) distributions generated by 5000 simulations of the simple cell diffusion model (Results subsection 3) from a single point to the tumor boundary with (red) and without (teal) a fence. The initial point is directly above (two lattice spacings away) or 20 *µm* and centered on the fence. The green line corresponds to the analytic solution for the FPT distribution of the system without a fence. When a cell is centered directly above a fence, it has lower probability of reaching the tumor boundary in a given time *T*, and it takes longer on average to reach the boundary. **(c)** Plot of tumor access ratio for different fence lengths at various times. Dots correspond to tumor access ratio, *R*_*b*_(*T*), calculated from simulations of the simple fence model. b. The ratio decreases from 1 (full access) with no fence to 0 (no access) when the fence covers the entire tumor boundary. The black curve is the tumor access ratio calculated for *T*=2.5 days for multiple fence lengths as described in (d). **(d)** The number of cell trajectories which reach the tumor boundary within *T* after reflecting off the fence at least once (red) is obtained from the simulation. The number of trajectories which reach the boundary directly (green) is calculated analytically (see text). The black curve in (c) is obtained by adding the contributions of these two types of trajectories. **(e)** The simulation box with an added chemokine gradient. Starting positions (not shown) are identical to those in (a). **(f)** The tumor access ratio calculated using the simple chemotaxis model plotted for varying fence length and time periods for the fence which induces cell chemotaxis. Cells in the region y ≤ 40 *µm* (see (e)) drift towards the tumor boundary at a constant speed determined by the length of the fence. Outside this region, the cells diffuse. The tumor access ratio is unity with no fence as there is no chemotaxis with no fence present. With increasing fence length, as chemotaxis drift rates increase, the metric first increases but then decreases as the fence begins to cover more of the boundary decreasing physical access.

A tumor access ratio of unity implies the fence does not affect the tumor access and the ratio is reduced as the fence increasingly hinders the access of the motile non-tumor cells to tumor cells. We computed the ratio using analytical calculations and stochastic simulations (see Materials and Methods). Our calculations showed that for a fixed fence of length of 70 *µm*, which is a common fence length found in the IMC data, motile cell access to the tumor is reduced by over 10% over 2.5 days (**Figure 4c**). In the ABM described earlier (**Figure 3**), the motile cells are cytotoxic CD8+ T cells. The 10% decrease in the tumor access ratio in the simple diffusive model due to fencing is comparable to the average 7% increase in the average cancer cell population in the larger ABM. The tumor access ratio also depends on the simulation time, *T*. The fence remains fixed during the simulation, thus different values of *T* probe the effects of fences with different lifetimes τ_fence_ ≈ T. We evaluate *R*_*b*_(*T*) for a range of *T* up to the typical timescale over which a tumor cell replicates once (∼2.5 days).

The full fencing metric is shown in Figure 4c. *R*_*b*_(*T*) is close to 1 when *b* << L and goes to zero as *b* approaches L. We find that, as *T* is reduced from 2.5 days, the tumor access ratio collapses onto a diagonal line displaying a linear relationship between the fence length and the tumor access ratio (**Figure 4c**). We can explain this as follows: each diffusing cell generates a trajectory through the lattice in time and each of these cell trajectories can be separated into three groups. One subset consisting of *N*_*nr*_ trajectories that never reflect off the fence but reach the tumor boundary in *T*, another subset of *N*_*r*_ trajectories that reflect off the fence of length *b* at least once before reaching the tumor boundary within *T*, and the remainder of the trajectories that never reach the tumor boundary (**Figure 4d**). The tumor access ratio is given by 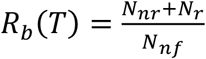 where *N*_*nf*_ is the number of cells that reach the tumor boundary when there is no fence. The first term, 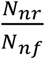, can be calculated analytically and is given by 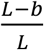. This corresponds to the linear dependence in Figure 4c. We compute the second term in 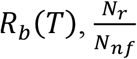, numerically as explained in the Materials and Methods which leads to the full expression of *R*_*b*_(*T*) seen in Figure 4c. When *T* is small, it is less likely for trajectories to reflect off the fence and subsequently reach the tumor boundary within *T*; i.e., *N*_*r*_ ≪ *N*_*nr*_ and *R*_*b*_(*T*) becomes linearly dependent on *b*.

The full nonlinear dependence of *R*_*b*_(*T*) on the fence length arises from the trajectories that reflect off the fence at least once then reach the tumor boundary. To uncover how fences affect the tumor access ratio further, we investigated how cells which reflect off of the fence contribute to the FPT distribution of the cells averaged over all the initial positions used to calculate *R*_*b*_(*T*). For a 90 *µm* fence, the cells which never reflect off the fence but reach the tumor boundary with *T* = 2.5 days, *N*_*nr*_, represent 69% of all trajectories which reach the tumor boundary with *T*. The other 31% of trajectories which contribute to the FPT reflect off the fence at least once. Generally, the higher the number of reflections off of the fence the smaller the number of cells which reach the tumor boundary after that number of reflections. In fact, we found that the fraction of all trajectories which reach the tumor boundary within time *T* which reflect off of the fence *n* times decreases exponentially with *n*. We have presented a quantitative characterization of the access of cytotoxic cells to cancer cells that determines the immune response for different values of the fence length and different durations in a simple geometry.

Next we investigated how, in some cases, fencing clusters can increase immune control of cancer cells in the TME. In our analysis we found that responders to therapy have on average greater formation of fencing clusters by TCF1+ CD4+ T cells and CD4+ T cells in TNBC and HNC respectively (**Figures 2a,b**). As some immune cells such as CD4+ T cells can recruit cytotoxic immune cells using chemokines, fences formed by these cells may enhance immune response. They could introduce a gradient of chemokines such as CCL3/4 and attract cytotoxic immune cells toward the tumor boundary (19). To investigate this, we calculated the increase of the tumor access ratio with the addition of cytotoxic cell chemotaxis to the simple diffusion model from earlier.

To do this, we first modified our original simplified fencing model adding a region in which the cytotoxic cell executes biased diffusion toward the tumor boundary as a simple method to account for chemotaxis (**Figure 4e**). The constant chemotactic drift rate of the motile cell increases with the fence length. This means that as the fence increases in size, the motile cell’s access to the tumor boundary is augmented with an increased drift rate but is diminished by the physical blockage of the tumor boundary. We calculate the tumor access ratio comparing the system with the fence generating chemotaxis to a system with no fence and therefore no chemotaxis (**Figure 4f**). With no chemotactic fence in place, the tumor access ratio is 1. As the fence is added and increases in length, the ratio first increases due to the effect of the drift. Ultimately, the ratio must decrease to 0 as the fence length spans the entirety of the system width. With the addition of chemotaxis-inducing fences, with *T* = 12*h*, the tumor access ratio reaches roughly 2 demonstrating an increase in cytotoxic cell access to the tumor boundary. This describes a mechanism by which tumor fencing benefits immune access and response in the TME.

## Discussion

The TMEs in solid tumors shows spatially heterogeneous organization of tumor and non-tumor cells. Spatial patterns that persist across patients and regulate tumor progression are being widely investigated at the single cell level. Here, instead of pursuing a data driven discovery of spatial patterns associated with patient prognosis (6-9), we focused on investigating the presence of a spatial pattern with simple geometric structure described by the fencing of tumor cells by diverse non-tumor (immune and non-immune) in the TME which can potentially mechanistically regulate tumor growth. We found the statistically significant occurrence of fencing clusters formed by a variety of immune cells such as CD4+/CD8+ T cell subtypes, B cells, NK cells, macrophages, granulocytes, and non-immune cells such as fibroblasts in the TME of lung, HNC, breast, brain, skin, and colorectal cancers. The frequency of occurrence of the fences depended on the cell type and the cancer type. For example, B cells formed fencing structures in over 75% of the 416 patients whereas Tregs formed fences in < 20% of the patients in the same cohort. Given the common presence of these structures across solid tumors it is possible that these structures, at the tumor-immune boundaries, arise due to some common interplay between cells in the TME. The local environment at the tumor-immune boundary could support increased proliferation of immune and non-immune cells (e.g., fibroblasts) which could potentially give rise to fencing clusters of the non-tumor cells. Previous investigations have shown that tumor cells are more transcriptionally active and there is increased localized production of IFNγ at the tumor immune boundary(13). Fencing clusters could also be generated through chemotactic aggregation as has been shown in ex vivo tumoroid models. Naïve CD8+ T cells which reach a tumoroid target chemotactically attract more CD8+ T cells in a feedback loop generating clusters of CD8+ T cells on the tumoroid boundary (20).We investigated association of fencing structures with patient response and survivability by developing a fencing participation metric, FPM. The FPM quantifies the participation of specific phenotypes of non-tumor cells to the fencing structures, i.e., the larger the value of FPM, the greater the fraction of the total number of a non-tumor cell type participates in forming the fencing structures. Note that a high FPM of some cell phenotype does not suggest that there is elevated contact between cells of that phenotype and cancer cells as the definition of a fencing cluster only requires contact of the cluster with a single cancer cell. Hence, an elevated FPM of cytotoxic CD8+ T cells does not imply elevated killing of cancer cells. The FPM for several non-tumor cell types showed statistically significant differences across patients responding or not responding to therapy. In TNBC, we found that non-responders to therapy (chemotherapy or ICI+chemotherapy) have on average a higher GzMB+CD8+ T cell FPM. Wang et al.(8, 21) analyzed the same dataset and found that the number of cancer cells in contact with the average GzMB+ CD8+ T cell is higher in responders as compared to non-responders to therapy. This may be explained by singlets and doublets of GZMB+ CD8+ T cells in contact with cancer cells which do not contribute to the FPM. For HNC, we found that higher values of FPM for CD8+ T cells, macrophages, and lymph vessels are associated with responders and increased survivability. The formation of fencing structures could lead to an enhanced immune response on the tumor immune boundary. For instance, non-tumor cells in fencing structures could generate chemokines or cytokines at the tumor boundary. We also found that increased FPM for M1-like microglia and non-classical monocytes in brain metastasis than primary brain tumors (glioma). There were several cancers where the FPM did not correspond to any statistically significant differences in patient outcomes. In lung cancer (lung adenocarcinoma), the FPM for B cells did not show any difference between responders and non-responders, whereas Sorin et al.(22), using the same dataset showed increased B cells in the cellular neighborhood is associated with greater survival. However, Sorin et al. (22) also found that, in lung cancer, cellular neighborhoods at the tumor boundaries, where fences are likely to form, are not associated with the survivability in a statistically significant manner. Therefore, in a range of solid cancers the spatial statistics of non-tumor cells (e.g., fencing structures) in physical contact with tumor cells appear to be relevant for tumor progression and patient prognosis.

Finally, we theoretically explored the dynamics of fencing and its effects on tumor progression. Using an ABM we developed previously to describe ICI response in melanoma patients(3), we showed that the average cancer cell population growth is increased by the presence of fixed fences due to reduced access of cytotoxic cells to cancer cells. We characterized this mechanism with a simple computational model which displayed that cytotoxic cell access to cancer cells is decreased by fences. Such a variation in cytotoxic cell access may play a role in the progression of patient TME which leads to response or non-response to therapy. We also constructed a simple computational model including chemotaxis to explore possible mechanisms by which fencing structure may benefit cytotoxic cell tumor access. This simple model showed that cytotoxic cell access to cancer cells may be augmented by chemotaxis driven by cells in a fencing structure. We have highlighted and investigated two mechanisms by which fences affect the progression of the tumor in silico. Ultimately, our models provide a framework with which we can draw conclusions about the effects of fencing as our understanding of physical and biochemical interplay between cells in the TME advances.

### Limitations of analysis

While the effects of fencing are not small, we do not expect fencing to be sole determinate of tumor clinical outcome. Several factors including tumor regions not considered in the TMA, other clinical confounding factors such as the age, co-morbidity factors can affect clinical outcomes. In addition, some of our datasets contained relatively small number of patients (e.g., CRC) which may preclude the determination of fencing occurrence by rare cell types. The tumors are three dimensional structures and fencing clusters found in two dimensional TMAs may underestimate those clusters formed in three dimensions(23). These limitations specific to the datasets can be overcome as the scientific community produces more and larger spatial tumor-imaging datasets. Experimental techniques which enable the imaging of many different cell phenotypes across entire tumor cross sections as well as in three-dimensions have recently been developed (15, 24, 25). In our simple computational models, we chose to simulate linear fences on linear tumor boundaries though fences and tumor boundaries manifest in a diverse array of shapes. Though the simple setup is expected to recapitulate some of important effects of fences on immune cell access to cancer cells, the quantitative aspects of the results can be modified by the detailed shape and composition (e.g., tumor vasculature(26)) of the tumor boundaries, and the factors such as cytokines, different immune cell types not considered in the simple models. Our study of immune cell access to cancer cells in simple fencing models can be extended to real tumor structures utilizing fence and tumor cell distributions taken from data.

## Materials and Methods

### Fencing existence calculation details

For the calculation of the probability of the occurrence of fencing clusters made of cell phenotype *g* fencing clusters in slide data, we constrained our calculations to only the subset of patient slides with at least 50 cells of a given phenotype *g* and 50 cancer cells. As mentioned in Result 1, we generated for each slide a p-value corresponding to the probability of the slide displaying at least as many *g*-cells in fencing clusters as compared to randomized data. We randomize the patient slides by sequentially choosing each individual non-cancer cell A and randomly swapping its position with another non-cancer cell B. Once cell A has been swapped, no other cell may swap with it. These swaps are performed until no non-cancer cells may be swapped. This randomization maintains the spatial structure of the cells in the slide. For each randomized slide, we find the number of *g*-cells which participate in fencing clusters. We generate over 1000 randomized slides and calculate the fraction of these randomized slides which have at least as many *g*-cells in fencing clusters as in the original patient slide. This fraction is an estimate of the probability that the number of *g*-cells participating in fencing clusters in the data is more than expected from randomly distributed non-cancer cells while preserving the slide’s spatial structure. This probability is a p-value.

We calculate this p-value for each patient slide forming a distribution of p-values. Using the Benjamini-Hochberg procedure to control the false discovery rate at level *p* = 0.05, we find the fraction of slides which display statistically significant tumor fencing.

For the lung cancer, HNC and brain cancer datasets, 10,000 randomized samples were generated per patient slide and per cell type to calculate both the p-value and the fencing participation metric. The number of random configurations used for the same calculations for the melanoma, breast cancer and colorectal cancer datasets were 6,000, 5,600 and 1,000 respectively. For each dataset, the number of randomizations performed per slide is at or above a chosen value such that the false discovery rate estimation could be controlled at at least p=0.05.

### Fencing participation metric for response prediction

We calculate the fencing participation metric (FPM) of phenotype *g*-cells for a patient slide with 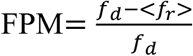, where *f* is the number of *g-*cells found in fencing clusters in the patient slide and < *f*_*r*_ > is the average number of *g-*cells found in fencing clusters in the set of randomized slides. The set of randomized slides are obtained using the same process used to generate randomized slides for the p-value calculation in the last section. We then map any negative FPM values to 0 to represent no participation in fencing structures as compared to randomized cell distributions. To compare the average FPMs between patient cohorts corresponding to different clinical outcomes, we utilize the t-test. For the datasets in which patient survival data was available, we also compared the survival curves of patient groups with high versus low cell phenotype FPM for each non-cancer cell phenotype. To separate patients into two groups of high and low FPM of cell phenotype *g*, we first found the average *g*-cell FPM across all patients. We then split the patients into those with equal or greater FPM (high) or smaller FPM (low) as compared to the average FPM across all patients. All survival curve comparisons shown differ with p<0.05 by log-rank test.

The FPM of a cell phenotype *g* in a patient slide has dependence on the density of cell type *g* as well as the densities of other cells in the patient slide. For example, when the system is filled with only cancer cells and *g*-cells (the density of other non-cancer cell types is 0), randomly generated configurations of *g*-cells are assured to reproduce any fencing structures found in the data. This means that the FPM will be 0 denoting that there is no evidence of fencing greater than expected from a random distribution of non-cancer cells. We also expect that, with a constant density of *g*-cells forming some fencing clusters, as the density of other cells which are not forming fencing structures increases (total non-cancer cell density increases), the FPM increases.

It has been found that patient tumor progression correlates with the increased or decreased densities of particular cell phenotypes in the TME (5). As the FPM depends on the density of the cells in the patient slide, a natural question is whether the FPM correlates with response only because of its dependence on cell density. We use the fencing and density of GZMB+ CD8+ T cells in TNBC as the central point of our analysis on the interdependencies of cell fencing with cell density for predicting patient response.

First, we calculate the density of GZMB+ CD8+ T cells, *σ*_*A*_, in each patient slide which has at least 50 GZMB+ CD8+ T cells and 50 cancer cells (so we may calculate the FPM as well). Each patient slide was assigned either a responder or non-responder to therapy status in the TNBC study. We assign responders a value of 1 and non-responders a value of 0 for the purpose of calculating correlations between density or FPM and response to therapy. GZMB+ CD8+ T cell density is positively correlated with response to therapy in TNBC with a Pearson correlation of 0.18. The GZMB+ CD8+ T cell FPM, *M*, calculated for the same patient slides correlates with response to therapy with a Pearson correlation of 0.14. To make sure that the FPM does not only correlate with response to therapy due to its dependency on cell density, we build a modified metric. We combine these two metrics into 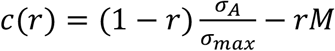, where *σ*_*max*_ is the maximum density of GZMB+ CD8+ T cells across all slides so that 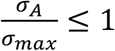. When r is 0, *c* has the same correlation with response as the cell density. When r is 1 it has the same correlation with response as the FPM. Because *M* ranges from 0 to 1, *c* ranges from −1 to 1. Plotting the Pearson correlation of *c* with patient response against r, we find if the addition of the FPM increases the correlation with response. If the FPM provides the same information on patient response to therapy as the cell density, we expect the correlation of *c* with response to monotonically decrease with increasing r. As we can see (**Figure S2a**), the correlation with response increases with the addition of the FPM up to the ratio of 0.2 after which it begins to descend to 0.14. For some cell types, the FPM has a stronger correlation with response as compared to the same cell’s density (**Figure S2b**). This shows that the cell FPM supplies information of response independent of the cell density.

### Simple fence computational model details

The setup of the simple computational model is as follows. We simulate the diffusive motion of a single cell on a two-dimensional lattice (lattice constant a=10 μm, corresponding to the cell size). Periodic boundary conditions are imposed along the width of the simulation box which is chosen to be roughly the average distance between fencing clusters in the TNBC data (230 μm) (**Figure 4)**. The length of the simulation box extends in the range of *y* ≥ 0. The lower boundary, *y* = 0 or the x-axis, is the upper edge of the tumor region and is the tumor boundary for the diffusing cell. The cell diffuses at a rate given by 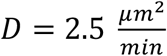 starting from its initial position. We place a fence of other cells with a specified length, *b*, just above and along the tumor boundary; it acts as a barrier reflecting the diffusing cell thus blocking access to the tumor boundary. We are interested in the ratio of the number of cells that reach the tumor boundary with the fence to that without, in a time *T*. We simulated 5000 samples and counted the fraction of samples in which the cell crossed the boundary in a time *T*, ranging from 0.5 to 2.5 days and for different fence lengths, including *b* = 0, i.e., no fence. The maximum value of *T* was chosen to match the time scale at which a cancer cell proliferates. The initial position of the cell was chosen to vary over a grid of points (see **Figure 4a**) chosen so that a cell has at least a 50% chance of reaching the boundary in 2.5 days when there is no fence. At a distance of 90 μm from the tumor boundary the cell has a 50% chance of reaching the boundary and so all the grid points were within 90 *µm*. The simulation box is restricted by *y* = 1500 *µm*, since any cell that reaches that distance has almost no chance of reaching the tumor boundary.

We can utilize the diffusive currents (the number of cells reaching the tumor boundary between times *t* and *t* + *dt* for a small time *dt*) into the entirety of the tumor boundary from the ensemble of initial conditions to calculate the tumor access ratio. It is defined in terms of the diffusive currents into the tumor boundary with a fence, *I*_*f*_(*t*), and without a fence, *I*_4*f*_(*t*), the ratio is defined

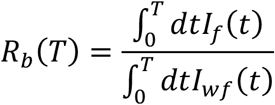

The denominator of this expression may be calculated analytically. First we calculate the exact solution for the first passage time distribution of a cytotoxic cell starting from a single starting position (at coordinates (*x*_6_, *y*_6_) = (0, *y*_6_), with arbitrary *y*_6_) with no fence. We start by constructing the probability density distribution *P*(*x, y, t*|0, *y*_6_, *t* = 0). Integrating this probability density over some area *A* yields the probability that the cell (started at (0, *y*_6_)) is found within that region at some time *t, Prob*(*cell is in A, t*) = ∬_*A*_ *P*(*x, y, t*|0, *y*_6_, *t* = 0)*dA*. Using the diffusive Green’s function (27), we superpose the probability distribution generated by a source at (*x, y*) = (0, *y*_6_) and a negative mirror source at (*x, y*) = (0, −*y*_6_) to satisfy the absorbing boundary condition at y = 0. Adding these, we construct the cell probability density function

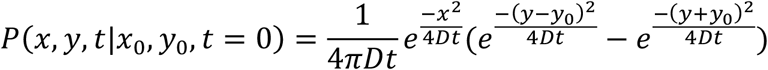

Going forward, when *P*(*x, y, t*) is written, is assumed conditional on starting at position (*x*_6_, *y*_6_) at *t* = 0. We integrate this function over the half plane (x > 0) to find the probability that the cell does not reach the x-axis by time t. The negative derivative of this integral with respect to time is the first passage time distribution *FPT*_4*f*_(*t*). Note that this first passage time is equivalent to the probability density current into the cancer region from a single initial condition, *I*_4*f*_(*t*|*y*_6_, *t* = 0). This current is independent of the starting position of the cell due to translational symmetry of the system along the x-axis.

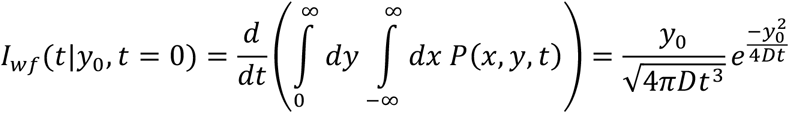

With this expression in hand, to produce the denominator, 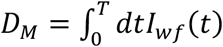, of the tumor access ratio, we average over all initial positions, *y*_*i*_, and integrate over the period, *T*

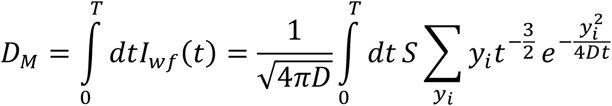

where *S* = 5000 is the number of samples simulated from each initial condition.

### Minimal model with chemotaxis

The simple computational chemotaxis model simulates the drift-diffusive motion of a single cell on a two-dimensional lattice (lattice constant a=10 μm, corresponding to the cell size) similar to that of the previous simple cell diffusion model. Periodic boundary conditions are imposed along the width of the simulation box which is chosen to be roughly the average distance between fencing clusters in the TNBC data (230 μm) (**Figure 4)**. The length of the simulation box extends in the range of *y* ≥ 0. The lower boundary, *y* = 0 or the x-axis, is the upper edge of the tumor region and is the tumor boundary for the diffusing cell. The cell diffuses at a rate given by 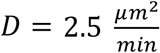. The cytokine gradient induced by the fence is included simply by the addition of a drift if the cells is within 50 *µm* from the tumor boundary, a typical chemokine decay distance (28). The drift which cytotoxic cells experience is directed towards the tumor boundary. The rate at which the cytotoxic cell chemotaxes (drifts) towards the tumor boundary linearly increases with the length of the CD4+ T cell fence from 0 to the maximum rate 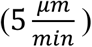 when the CD4+ T cell fence reaches 150 *µm*. Drift rates of this magnitude have been observed in CD8+ T cells (29). The drift rate is assumed constant with further increases in fence length past 150 *µm*. For the simple fence chemotaxis model, the same method of calculating the tumor access ratio in the diffusion model is used. It compares the number of motile cells which reach the tumor boundary within *T* with a fence to the number of motile cells which reach the tumor boundary within *T* without a fence and therefore without chemotactic drift.

### Modified Interacting Cell System (ICS) Model

The computational model applied to find the effect of fencing on tumor progression in silico is a modified version of the Interacting Cell System (ICS) model (3). The primary modifications to the ICS model are the following: i) We implant a few fences consisting of a new type of non-tumor cells to the initial configuration of the tumor and the immune (T-cells and macrophages) cells considered earlier. The non-tumor cells in the fence do not interact with tumor or T cells and macrophages; nor do they proliferate, die, or move during the simulation. ii) When activated CD8+ T cells are converted to T_ex_ cells they immediately removed from the system. This rule is implemented to make sure the effect of our implanted fencing barriers are not affected by fences formed by T_ex_ cells. The other details of the ICS model are similar to that described in ref. (3). Below we briefly describe main features of the modified ICS model.

The Interacting Cell System (ICS) model is a kinetic Monte Carlo simulation implemented on a 100×100 square lattice with spacing 10 *µm* representing the 1mm×1mm TMAs taken from each patient. We customized SPPARKS Kinetic Monte Carlo Simulator (distributed at spparks.github.io) code to build this agent-based model. The references supporting model rules and parameter values can be found in ref. (3). The physical extent of the cells constrains the number of cells in each chamber that corresponds to a 10 *µm* × 10 *µm* square. The model considers four types of cells in the system: tumor cells, activated CD8+ T cells, TAMs, and non-tumor cells in the implanted fences. Activated CD8+ T cells occupy a fourth of a chamber while tumor cells and macrophages each fill half a chamber. These occupation rules reflect the limits of occupation in the original IMC slides. A chamber with a fence cell is fully occupied. Two cells are considered in contact if they are in the same chamber or in adjacent chambers that share an edge.

The initial state of the system is obtained from the patient’s TMA by discretizing the cell positions in the IMC data for a patient TMA. Over time, cell positions and numbers change: First, tumor cells proliferate (rate *k*_*C*_) with the location of the daughter depending on the availability of space in progressively larger neighboring regions. The new cell is placed in the same chamber as the proliferating cell if space is available, if not then the surrounding 8 chambers, next the adjacent layer of 16 chambers and finally the third layer of 24 chambers are checked for available space. If space cannot be found in the 49-cell neighborhood, even by expelling CD8+ T cells, then proliferation does not occur. When multiple chambers are available in a layer, one is chosen randomly.

Activated CD8+ T cells proliferate (30) into the same chamber they occupy if the chamber is not already full. The rate at which activated CD8+ T cells proliferate is limited by CD8+ T cell competition for resources and space in the TME. We include this competition by enforcing a maximum number (or carrying capacity) of the CD8+ T cell population, *N*_*cc*_. Given the number of activated CD8+ T cells, 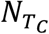, and exhausted CD8+ T cells, 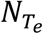, the total CD8+ T cell population is 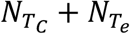. The carrying capacity is incorporated in the rate of proliferation and recruitment with a logistic term 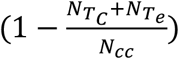 which is 1 when there are no CD8+ T cells and 0 when the population reaches the carrying capacity. The rates at which activated CD8+ T cells proliferate and recruit to the TME also depend on the number of recently killed cancer cells (31, 32). As the number of tumor cells killed in the previous 12 hours (effective immune system memory span, *T*_*m*_), *D*_*C*_, increases, the proliferation and recruitment rates grow until they reach their maximum values. We employ a Michaelis-Menten functional form to capture this saturation of the proliferation and recruitment rates at maximum values: 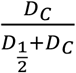 where 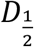 is the Michaelis constant, i.e., when 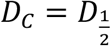, the rate is halved. The more tumor cell death, the greater the probability of neoantigens being created invigorating activated CD8+ T cell proliferation and recruitment by way of antigen-presenting cells (APCs). This constitutes a positive feedback for activated CD8+ T cell population. APCs are not explicitly modeled by our ICS model but their effects on CD8+ T cell proliferation and recruitment are included. Putting these together, we have

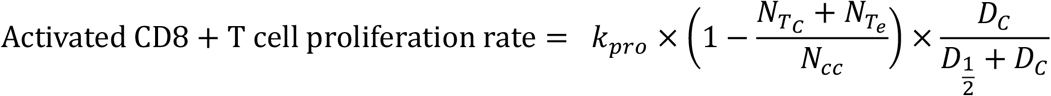

where *k*_*pro*_ is the maximum proliferation rate of activated CD8+ T cells.

Activated CD8+ T cells are recruited at a rate with the same dependencies on CD8+ T cell carrying capacity and killed tumor cells as the activated CD8+ T cell proliferation rate,

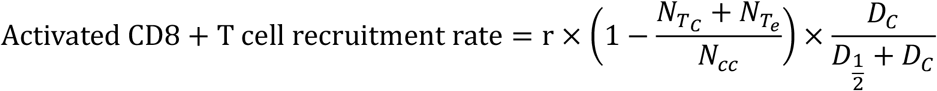

where *r* is the maximum recruitment rate of activated CD8+ T cells. When recruited, activated CD8+ T cells are randomly placed in a chamber with available space for the cell to occupy. TAMs are recruited at a constant rate, *r*_*M*_, and are similarly placed randomly in a lattice chamber with enough space to accommodate the TAM. TAMs die at a constant rate *γ*_*M*_.

In the model, activated CD8+ T cells are exhausted by tumor cells and TAMs with which they are in contact. Once exhausted, they are removed from the system. In vivo, tumor cells express PDL1 and PDL2 (33-35) and chronic PD1 activation in CD8+ T cells leads to exhaustion (36). TAMs express PDL1 and PDL2 in the TME (35) and may directly inhibit activated CD8+ T cells (37-40). In CD8+ T cells, PD1 and CTLA4 expression upregulates upon TCR engagement (41, 42). These interactions lead CD8+ T cells to lose cytotoxic and proliferative ability (43).

We account for cell motility in the TMA by allowing CD8+ T cells and TAMs to diffuse on the lattice. A motile cell may move into one of the four nearest neighbor chambers if there is sufficient space for the cell to occupy. The diffusion rate of CD8+ T cells is reduced when in contact with TAMs from *D*_*mot*_ by a factor of 33. This is a coarse-grained approximation to the myriad of complex processes that promote contact between CD8+ T and tumor cells. We apply periodic boundary conditions on our lattice which may be viewed as cell migration to and from other regions of the TME not shown in the data. When crossing a lattice boundary, CD8+ T cells have a 50% chance of being removed from the system.

### Data Availability

Data used to generate all figures may be found at https://doi.org/10.5281/zenodo.15817184. Further ICS model description, and instructions and code to build and execute ICS model simulations can be found at in our previous paper (3) and at https://github.com/gdag458/Melanoma_ICS. The patient datasets containing all patient IMC slides are made available in the public domain by the authors of the published manuscripts (6-8, 16-18).

## Acknowledgments

We thank the Abigail Wexner Research Institute at the Nationwide Children’s Hospital for funding.

## Supplementary Information for

**Table S1:** A table providing the cell phenotypes in the order (left to right) in which their fencing slide percentage appears in **Figure 1e**. For each cancer type-cell phenotype bin, the corresponding list of cell sub-phenotypes are listed in order of appearance from left to right in **Figure 1e**. The number of patient slides in which the fencing of the corresponding cell type was evaluated and the percentage which do display fencing significantly (p<0.05) is provided. Each cell phenotype given was defined by the authors of the corresponding studies.

|  | CD8+ T | CD4+ T | B | NK | APC | Granulocyte | Fibroblast | Other |
| --- | --- | --- | --- | --- | --- | --- | --- | --- |
| Lung Cancer | Cytotoxic CD8+ T (Tc) 50.87% of 401 slides | CD4+ Helper T (Th) 69.90% of 412 slides, Treg 17.87% of 263 slides | B 80.39% of 255 slides | NK 25.53% of 47 slides | Alt. Macrophage 65.34% of 326 slides, Cl. Macrophage 29.09% of 416 slides, Cl. Monocyte 53.18% of 299 slides, DC 57.14% of 14 slides, Intermediate Monocyte 32.41% of 108 slides, Non-classical Monocyte 38.26% of 230 slides | Neutrophil 57.45% of 322 slides |  | Endothelial 77.83% of 415 slides, Mast 40.11% of 177 slides, T other 8.52% of 317 slides |
| Brain Cancer | Cytotoxic CD8+ T (Tc) 50% of 118 slides | CD4+ Helper T (Th) 66.07% of 112 slides, Treg 0% of 4 slides | B 40.91% of 44 slides |  | DC 43.24% of 37 slides, Classical Monocyte 25.51% of 98 slides, Non-classical Monocyte 14.52% of 124, Intermediate Monocyte 32.26% of 62 slides, M1-like MDM 54.16% of 373 slides, M2-like MDM 49.70% of 328 slides, M1-like Microglia 23.17% of 259 slides, M2-like Microglia 21.81% of 188 slides | Neutrophil 64.37% of 87 slides |  | T other 30.30% of 66 slides, Mast 75% of 12 slides, Astrocyte 39.13% of 345 slides, Endothelial 80.41% of 342 slides |
| Melanoma | Naive CD8+ T 30% of 10 slides, Activated CD8+ T (Tc ae) 80% of 15 slides, Ex. CD8+ T 71.43% of 14 slides | Naive Helper CD4+ T 50% of 4 slides, Activated Helper CD4+ T (Th ae) 76.92% of 13 slides, Treg 25% of 8 slides | B 75% of 4 slides |  | Macrophages & Monocytes 40% of 25 slides |  |  | CD31+ 76% of 25 slides |
| Head and Neck Cancer | CD8+ T 74.36% of 39 slides | CD4+ T 81.82% of 308 slides | B 85.71% of 7 slides |  | Macrophage 88.96% of 308 slides, APC 73.33% of 90 slides | Granulocyte 76.24% of 282 slides | Stroma/Fibroblast 63.64% of 308 slides | Naive Immune 69,18% of 292 slides, Lymph vessel 60.71% of 112, Vessel 91.78% of 304 slides |
| Colorectal Cancer | CD8+ T 85.71% of 7 slide | CD4+ T 100% of 7 slides, Treg 100% of 7 slides | B 100% of 5 slides |  | Macrophages & Monocytes 100% of 8 slides, IL17a+ Macrophage 100% of 7 slides |  | Fibroblast 75% of 8 slides | T 100% of 1 slide, Neutro & NK 100% of 6 slides, Fibro & Mono 100% of 8 slides |
| Triple Negative Breast Cancer | CD8+ T 16.78% of 447 slides, TCF1+ CD8+ T 14.29% of 203 slides, Gzmb+ CD8+ T 30.14% of 282 slides, Ex. CD8+ T 37.42% of 310 slides | TCF1+ CD4+ T 32.48% of 545 slides, PD1+ CD4+ T 26.27% of 354 slides, Treg 4.41% of 476 slides | CD20+ B 45.10% of 204 slides, CD79a+ Plasma 47.23% of 451 slides | CD56+ NK 13.43% of 67 slides | DC 18.58% of 479 slides, M2 Macrophage 20.06% of 658 slides, PDL1+ APC 47% of 200 slides, PDL1+ IDO+ APC 66.87% of 166 slides | Neutrophil 57.03% of 128 slides | Fibroblast 5.84% of 1199 slides, Myofibroblasts 31.52% of 330 slides | PDPN+ Stromal 25% of 544 slides, CA9+ 57.14% of 49 slides |

**Figure S1.**
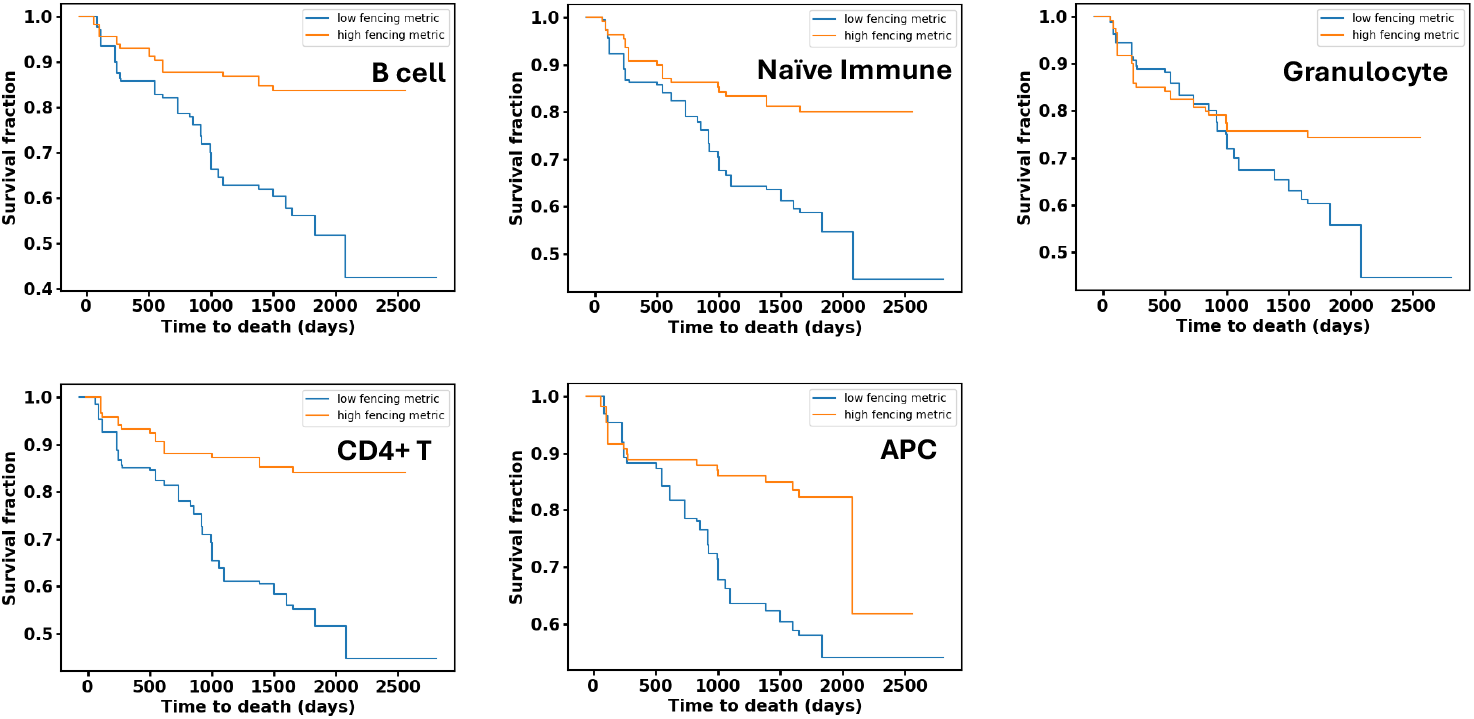
Fencing participation metric for different cell types is associated with survival of patients in HNC. For HNC, we find that the survival of patients with high (above average) and low (below average) fencing participation metric calculated for certain cell types differ significantly (using the log-rank test, p<0.05). Here we show the survival curves for those cell types. Each curve is generated by the survival data yielded from over 100 patients.

**Figure S2.**
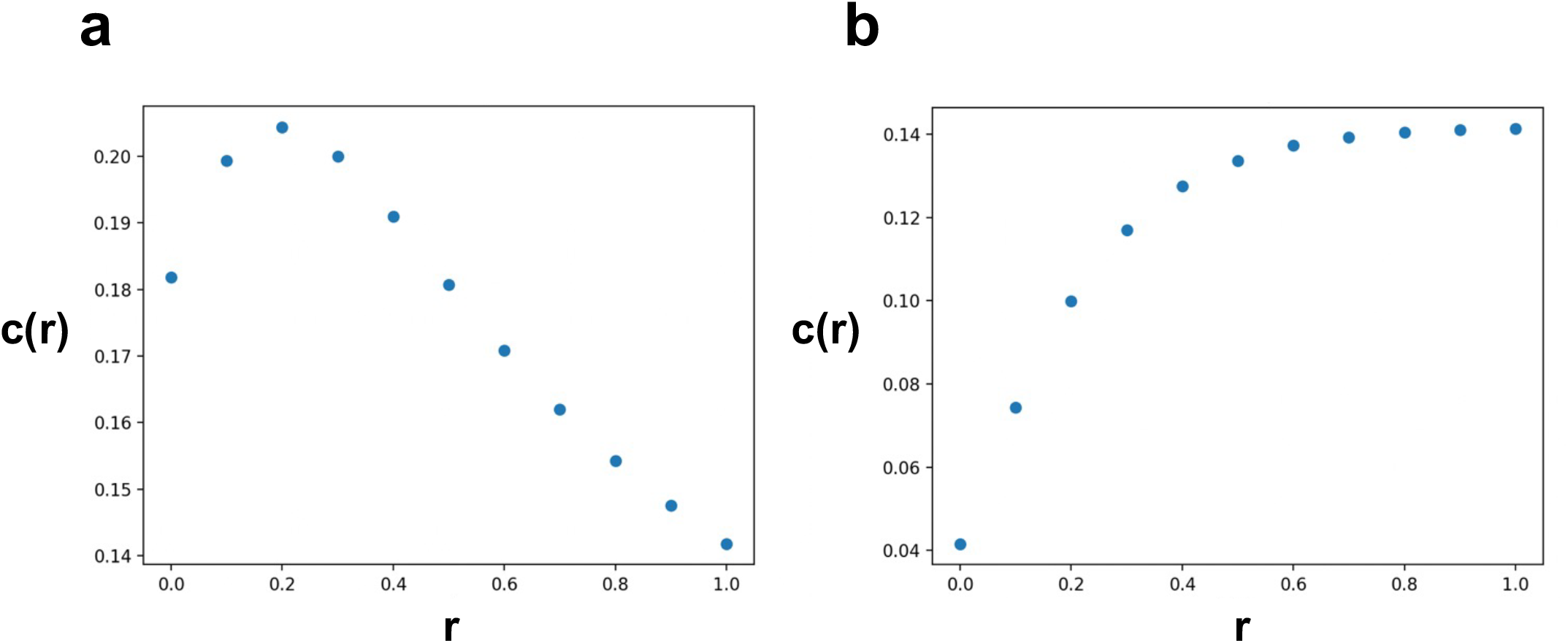
Characterizing the relation between cell density and the fencing participation metric for TNBC. The correlation of 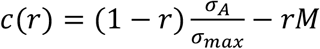 with patient response in TNBC plotted against the ratio r for **(a)** GZMB+ CD8+ T cells and **(b)** CD4+ PD1+ T cells. At r=0, we find the correlation of cell density with response. In **(a)**, introducing the contribution of the fencing participation metric to *c*(*r*) by increasing r, increases the correlation with response before it decreases. In **(b)**, the fencing participation metric has a stronger correlation with response than the density for CD4+ PD1+ T cells

